# Novel FKBP12 ligand promotes functional improvement in SOD1^G93A^ ALS mice

**DOI:** 10.1101/2024.01.17.576010

**Authors:** Laura Moreno-Martinez, Núria Gaja-Capdevila, Laura Mosqueira-Martín, Mireia Herrando-Grabulosa, Klaudia Gonzalez-Imaz, Ana C. Calvo, Maialen Sagartzazu-Aizpurua, Leticia Moreno-García, Jose Manuel Fuentes, Abraham Acevedo-Arozena, Jesús María Aizpurua, José Ignacio Miranda, Adolfo López de Munain, Ainara Vallejo-Illarramendi, Xavier Navarro, Rosario Osta, Francisco Javier Gil-Bea

**Affiliations:** Centro de Investigación Biomédica en Red Sobre Enfermedades Neurodegenerativas (CIBERNED), 28031 Madrid, Spain; LAGENBIO, Faculty of Veterinary, University of Zaragoza, Miguel Servet 177, 50013 Zaragoza, Spain; Aragón Health Research Institute (IIS Aragón), Biomedical Research Centre of Aragón (CIBA), 50009 Zaragoza, Spain; AgriFood Institute of Aragon-IA2 (UNIZAR-CITA), 50013 Zaragoza, Spain; Department of Cell Biology, Physiology and Immunology, Institute of Neurosciences, Universitat Autònoma de Barcelona, 01893 Bellaterra, Spain; Group of Neurosciences, Departments of Pediatrics and Neuroscience, Faculty of Medicine and Nursing, University of Basque Country (UPV/EHU), 20014 San Sebastian, Spain; Department of Neuroscience, Biodonostia Health Research Institute (IIS BioGipuzkoa), 20014 San Sebastian, Spain; Department of Organic Chemistry-I, Korta Research Center, University of the Basque Country (UPV/EHU), 20018 San Sebastian, Spain; Department of Biochemistry and Molecular Biology and Genetics, Faculty of Nursing and Occupational Therapy, University of Extremadura, 10003 Cáceres, Spain; Instituto de Investigación Biosanitaria de Extremadura (INUBE), Cáceres, Spain; Research Unit, Canarias University Hospital, ITB-ULL, 38320 Tenerife, Spain; Miramoon Pharma, S.L., 20014 San Sebastian, Spain; Donostia University Hospital, 20014 San Sebastian, Spain; IKERBASQUE Basque Foundation for Science, 48009 Bilbao, Spain; Department of Health Sciences, Public University of Navarra, 31008 Pamplona, Spain

## Abstract

Amyotrophic lateral sclerosis (ALS) is a devastating neurodegenerative disease with limited treatment options. ALS pathogenesis involves intricate processes within motor neurons (MNs), characterized by dysregulated Ca^2+^ influx and buffering in early ALS-affected MNs. This study proposes the modulation of ryanodine receptors (RyRs), key mediators of intracellular Ca^2+^, as a therapeutic target. A novel class of novel FKBP12 ligands that show activity as cytosolic calcium modulators through stabilizing RyR channel activity, were tested in the SOD1^G93A^ mouse model of ALS. Different outcomes were used to assess treatment efficacy including electrophysiology, histopathology, neuromuscular function, and survival. Among the novel FKBP12 ligands, MP-010 was chosen for its central nervous system availability. Chronic administration of MP-010 to SOD1^G93A^ mice produced a dose-dependent preservation of motor nerve conduction, with the 61 mg/kg dose significantly delaying the onset of motor impairment. This was accompanied by improved motor coordination, increased innervated endplates, and significant preservation of MNs in the spinal cord of treated mice. Notably, MP-010 treatment significantly extended lifespan by an average of 10 days compared to vehicle. In conclusion, FKBP12 ligands, particularly MP-010, exhibit promising neuroprotective effects in ALS, highlighting their potential as novel therapeutic agents. Further investigations into the molecular mechanisms and clinical translatability of these compounds are needed for their application in ALS treatment.

## INTRODUCTION

Amyotrophic lateral sclerosis (ALS) is a devastating neurodegenerative disease that targets both upper and lower motor neurons within the brain and spinal cord. Tragically, ALS is characterized by a brief survival span of 2-3 years, with only 10-20% of patients surpassing a decade of life post-diagnosis. Familial cases, attributable to genetic mutations, account for 5-10% of all ALS cases, with the rest classified as sporadic (Mead et al., 2023). Despite intense ongoing research, current medications such as Riluzole, Edaravone, Relyvrio, and Qalsody provide only a modest delay in disease progression. The pathogenesis of ALS is a complex interplay of multiple processes occurring within motor neurons (MNs), encompassing amongst other proposed mechanism: excitotoxicity, endoplasmic reticulum (ER) stress, mitochondrial dysfunction, oxidative stress, RNA processing aberrations, protein misfolding, endosomal trafficking disruptions, impaired axonal transport or neuroinflammation (Hardiman et al., 2017; Zhou and Xu, 2023). Computational models have underscored the significance of disturbances in calcium (Ca^2+^) homeostasis and energy imbalances as pivotal factors predicting neuronal dysfunction and degeneration in ALS (LeMasson et al., 2014).

One distinguishing feature in early ALS-affected MNs is the excessive influx of Ca^2+^ ions due to high-frequency rhythmic activity and an increased expression of Ca^2+^-permeable AMPA receptors (Guatteo et al., 2007; Bursch et al., 2019). These MNs exhibit low expression of Ca^2+^ buffering proteins like parvalbumin and calbindin (Ince et al., 1993; Alexianu et al., 1994). Overexpression of these proteins has been shown to confer resistance to ALS-induced toxicity in MNs (Ho et al., 1996). Conversely, MNs that exhibit resistance to ALS degeneration, such as those of oculomotor and Onuf’s nuclei, possess higher levels of Ca^2+^ buffering proteins (Elliott and Snider, 1995; Vanselow and Keller, 2000). Notably, ALS-associated mutations, including those in *VAPB*, *SIGMAR1* and *SOD1* can potentiate Ca^2+^ deregulation and heighten vulnerability to the effects of Ca^2+^ influx (De vos et al., 2012; Bernard-Marissal et al., 2015; Leal and Gomes, 2015; Petrozziello et al., 2022). These findings emphasize the inherent susceptibility of MNs to intracellular Ca^2+^ overload as a significant risk factor for degeneration (Appel et al., 2001).

Mitochondria play a pivotal role in regulating intracellular Ca^2+^ levels through the mitochondrial uniporter, particularly crucial in MNs with low intrinsic cytosolic Ca^2+^ buffering capacity. In these vulnerable MNs, mitochondria are responsible for absorbing over 50% of intracellular Ca^2+^ increases, setting them apart from other cell types. Consequently, the pronounced demand placed on specialized Ca^2+^ storage organelles makes MNs even more susceptible to Ca^2+^ imbalances (Bergmann and Keller, 2004; Jaiswal, 2013), especially in ALS where mitochondria are frequently damaged (Zhou and Xu, 2023). Indeed, impaired mitochondrial Ca^2+^ buffering in MNs derived from patient-induced pluripotent stem cells carrying mutations in *TARDBP* or *C9orf72* has been reported (Dafinca et al., 2020).

Apart from excessive Ca^2+^ influx and diminished Ca^2+^ buffering capacity, augmented Ca^2+^ leakage from ER stores in MNs can further contribute to cytosolic Ca^2+^ overload. Ryanodine receptors (RyR) are tetrameric channels that play a crucial role in mediating Ca^2+^-induced Ca^2+^ release from the ER, play a crucial role in this process (Fabiato, 1983; McPhersonx et al., 1991). RyR channels are activated by Ca^2+^ and can also be triggered by the oxidation of redox-sensing thiols by reactive oxygen species (ROS), creating a positive feedback loop exacerbating pathological cytoplasmic Ca^2+^ overload (Zima and Mazurek, 2016). The endogenous ligand of RyR, FKBP12 (calstabin), normally stabilizes RyR, preventing Ca^2+^ leakage from the ER. Reduced FKBP12 levels in MNs of ALS patients underscore the importance of maintaining equilibrium between FKBP12 and RyR in neurodegeneration (Kihira et al., 2005).

Overall, this cumulative evidence suggests the prospect of RyRs as a potential therapeutic targets for ameliorating intracellular Ca^2+^ dysregulation and forestalling Ca^2+^-induced neurodegeneration within susceptible neurons and neuromuscular units in ALS. In this context, we introduce a novel class of triazole molecules, empirically validated as FKBP12 ligands to facilitate FKBP12-RyR binding. This binding capability serves to stabilize the Ca^2+^ flux mediated by RyR, as demonstrated in our previous works (Aizpurua et al., 2021; Passannante et al., 2023). In this study, we introduce MP compounds, novel FKBP12 ligands that exhibit cytosolic calcium modulating activity, with the potential to be used for the treatment of ALS. This proposal is based on the encouraging results observed in transgenic SOD1^G93A^ mice, a mouse model that recapitulates the key clinical, electrophysiological, and histopathological features of the disease. (Turner and Talbot, 2008; Mancuso et al., 2011).

## METHODS

### Preparation of MP compounds

Compounds MP-001 and MP-002, strategically designed to target the interaction site of FKBP12/RyR1, were synthesized using CuAAC “click chemistry” methodologies. Specifically, aryl propargyl sulfides and methyl azidoglycinate were utilized for MP-001, while 2-azidoethyl-N,N-dimethylammonium hydrochloride was employed for MP-002 synthesis (Aizpurua et al., 2021; Passannante et al., 2023). Compound MP-010 was prepared by adding phthaloyl peroxide (Gan et al., 2017) to a solution of MP-002, followed by stirring at room temperature, basification with 7 N NH_3_ in MeOH, and evaporation under reduced pressure. The resulting crude product underwent purification through column chromatography (silica gel; eluent: CH_2_C_l2_ / MeOH (7N NH_3_)). The yield of the purified white solid was 66%, with ^1^H and ^13^C NMR spectra confirming the chemical structure: ^1^H NMR (400 MHz, CD_3_OD) δ 7.81 (s, 1H), 7.49 (d, *J* = 8.8 Hz, 2H), 7.95 (d, *J* = 8.8 Hz, 2H), 4.51 (t, *J* = 6.3 Hz, 2H), 4.30 (dd, *J* = 13.7, 1.6 Hz, 2H), 3.86 (s, 3H), 2.82 (t, *J* = 6.3 Hz, 2H), 2.31 (s, 6H). ^13^C NMR (101 MHz, CD_3_OD) δ 164.1, 137.5, 133.5, 127.7, 126.7, 115.9, 59.5, 56.1, 53.6, 45.4. IR (cm^-1^): 1592, 1495, 1459, 1303, 1249, 1172, 1086, 1022, 829, 524.

To obtain aqueous 10 mM stock solutions of these compounds for pharmaceutical application, the free carboxylic acid of MP-001 or the tertiary amine moiety of MP-010 was neutralized to pH = 6.8 using 2 M NaOH and 2 M HCl, respectively. These neutralized solutions were further diluted with drinking water to achieve pharmaceutically acceptable concentrations of 1 mM or 2 mM prior to administration.

### Recordings of ER and cytoplasmic Ca^2+^ in HEK293 cells

HEK293 cells, stably and inducible-expressing WT and mutant R2474S RyR2, were generated as described (Murayama et al., 2015) and provided by Dr. Takashi Murayama (Juntendo University). Cells co-express R-CEPIA1er, a calcium-measuring organelle-entrapped protein indicator located at the ER, and they have an inducible expression for wild-type RyR2 and mutant RyR2 R2473S isoforms. 60.000 cells/well were seeded onto 96-well plates coated with 50 µg/ml poly-D-Lysine, as described previously. Cells were grown in 10% FBS, DMEM media, at 37°C and 5% CO2 for 24 hours. On day 2, RyR expression was induced with 2 µg/ml doxycycline. Induction was achieved after 24-28 hours, when calcium measurements were performed in HEPES-buffered Krebs solution (140 mM NaCl, 5 mM KCl, 1 mM MgCl2, 2 mM CaCl2, 11 mM D-(+)-Glucose, 5 mM HEPES, distilled H2O to 450 mL, pH 7.4). Fluorescent ratios (F/F0) were calculated as the ratio of the fluorescence intensities between the initial (F0) and the last (F) 100 seconds by R-CEPIA1er in the Glomax Discover Microplate Reader (Promega). Caffeine-induced Ca2+ transients were also measured with Fura2 AM at room temperature using an ECLIPSE Ti/L100 epifluorescence microscope (Nikon) equipped with a 20X S-Fluor objective, a lambda-DG4 illumination system, an Orca-Flash 2.8 camera (Hamamatsu) and NisElements-AR software.

### In vivo pharmacokinetics in mice

A bolus of 30 mg/kg of test compounds were administered via oral gavage to isoflurane-anesthetized control mice. Tissue samples, including muscle, brain, and serum, were collected at 2-, 4-, 6-, and 18-h post-administration, from three mice at each time point. Collected tissues were promptly preserved at -80°C until further processing. Frozen muscle and brain samples were pulverized in liquid nitrogen using a steel mortar immersed in dry ice. Approximately 100 mg of pulverized tissue and 100 µL of serum were then incubated with a 1% formic acid-acetonitrile solution, with equal volumes of tissue mass for muscle and brain, and three times for serum. Following a 90-s ultrasound bath for muscle and a 10-s bath for serum and brain samples, the extracts underwent centrifugation at 9600 g and 4°C for 5 minutes. The resulting supernatants were collected and stored at -80°C until further quantification. Quantitative determination of compounds in the biological samples was carried out using liquid chromatography-tandem mass spectrometry (LC-MS/MS) at the Research General Services SGIker of UPV/EHU (Vitoria-Gasteiz). The pharmacokinetic parameters following oral administration were calculated using the PKSolver add-in program (Zhang et al., 2010). Noncompartmental analysis was employed to compute the pharmacokinetic parameters of the parent compound. The terminal slope was automatically estimated using regression with the largest adjusted R^2^. The parameters inferred included the terminal half-life (t_1/2_), maximum concentration (C_max_), and the time taken to reach the maximum concentration (T_max_). Subsequently, the brain-to-serum (Cb:Cs) was calculated based on the obtained pharmacokinetic parameters (Shaffer, 2010).

### Animals and experimental design: Treatment of SOD1^G93A^ mice

We used the hybrid strain of SOD1^G93A^ mouse as an ALS model (B6SJL-Tg[SOD1-G93A]1Gur). Non-transgenic wild-type (WT) littermates were used as controls. The animals used were from colonies that had been bred in different animal facilities for years, as specified below. The transgenic offspring was identified by polymerase chain reaction (PCR) amplification of DNA extracted from the tail, to identify hemizygous transgenic mice and WT. The colony was maintained under stander conditions (food and water ad libitum, room temperature of 22 ± 2 °C and 12:12-h light–dark cycle) at the Animal Service of the UAB and at the animal facilities in Centro de Investigación Biomédica de Aragón University of Zaragoza. Mice were handled in accordance with the guidelines of the European Union Council (Directive 2010/63/EU) and Spanish regulations (RD53/2013) on the use of laboratory animals. All experimental procedures were approved by the Ethics Committee of the Universitat Autònoma de Barcelona (CEEAH-UAB: 2969 and 4273) and by the Ethic Committee for Animal Experiments from the University of Zaragoza (PI45/18, PI15/21, PI51/21, PI02/23). Humane endpoint criterium in the survival studies was considered when animals lost the righting reflex for longer than 30 s.

#### Trial 1: Effect of MP-010 on neuromuscular outcomes

This trial was conducted at the animal facility of the Universitat Autònoma de Barcelona using the colony of SOD1^G93A^ mice that has been bred over years by Dr. Navarro’s group. At 8 weeks of age (before drug treatment started), WT and SOD1^G93A^ mice were distributed among 6 experimental groups, according to their progenitors, sex, weight, and electrophysiology baseline values in balanced groups, receiving either vehicle or MP-010 at two different doses. The compound MP-010 was added to the drinking water at concentrations of 1 mM or 2 mM, resulting in average daily dosages of 61 mg/kg or 122 mg/kg, respectively. The mice were distributed in six groups for assessment of neuromuscular outcomes: 1) WT vehicle, this group consists of WT mice with no treatment (n=4/4; females/males); 2) WT MP-010 61 mg/kg (n=5/4); 3) WT MP-010 122 mg/kg (n=2/3); 4) SOD1^G93A^ vehicle (n=8/8); 5) SOD1^G93A^ MP-010 61 mg/kg (n=7/8); 6) SOD1^G93A^ MP-010 122 mg/kg (n=5/4). The treatment was given from the 8th until the 16th week of age and electrophysiological, behavioral and histological analyses were performed during this follow-up.

#### Trial 2: Effect of MP-010 on survival, behavioral tests, and genetic markers

This trial was conducted at the animal facility of the University of Zaragoza using the colony of SOD1^G93A^ mice that has been bred for years by Dr. Osta’s group. MP-010 was given to 21 SOD1^G93A^ mice (n=8/13 females/males) in the drinking water at a concentration of 1 mM, treating the mice at a dose of 61 mg/kg. Simultaneously, another group of 20 SOD1^G93A^ mice (n=9/11 females/males) was treated with vehicle as the control group. The treatment started at 8 weeks of age and was weekly renewed until each mouse reached its respective humane endpoint. Throughout the treatment, all mice underwent weekly weight measurements and motor behavioral tests, including rotarod and hanging-wire tests. All behavioral experiments in the three trials were conducted under blind conditions regarding the treatment.

#### Trial 3: Effect of MP-001 on survival, behavioral tests and genetic markers

This trial was conducted at the animal facility of the University of Zaragoza using the colony of SOD1^G93A^ mice that has been bred for years by Dr. Osta’s group. This experimental study included one group of 19 SOD1^G93A^ mice (n=9/10 females/males) to which compound MP-001 was administered in the drinking water at a concentration of 0.85 mM, resulting in 61 mg/kg dose, and another similar group of 17 SOD1^G93A^ mice (n=9/8 females/males) treated with vehicle as control. All mice underwent weekly weight measurements and behavioral tests, including rotarod and hanging-wire tests, and the survival time was monitored.

### Functional evaluation

#### Rotarod

The Rotarod test (ROTA-ROD/RS, LE8200, LETICA, Scientific Instruments; Panlab, Barcelona, Spain) was conducted on a weekly basis, spanning from 8/9 to 16 weeks of age or until mice were unable to perform the test in the case of mice from the survival experiment, to assess the motor coordination, muscle strength, and balance of the experimental animals. Specifically, the test involved measuring the duration for which each mouse could maintain its position on a rotating rod set at a constant speed of 14 revolutions per minute (rpm). Each animal underwent a series of five trials, and the longest latency achieved without falling was recorded. An arbitrary cut-off time of 180 seconds was applied in the assessment. The symptomatic disease onset for each mouse was determined as the first week when the mouse was unable to keep walking 180 seconds on the rod.

#### Hanging-wire test

Hanging-wire test was carried out in mice involved in the survival study to assess the muscular strength. Briefly, each mouse was placed on an inverted wire lid, and the time it took for the mouse to fall was recorded. Similar to the rotarod test, the mice had three attempts to remain on the wire for a maximum of 180 seconds per trial, and the longest latency was noted.

### Nerve conduction tests

Motor nerve conduction tests were performed by percutaneously sciatic nerve stimulation (with single square pulses of 20 μs duration) delivered through a pair of needle electrodes placed at the sciatic notch. Compound muscle action potentials (CMAP) were recorded from the tibialis anterior (TA) and plantar interossei (PL) muscles utilizing microneedle electrodes. Recorded potentials were amplified and displayed with settings adjusted to measure amplitude from baseline to the maximal negative peak, latency from the stimulus to the onset of the first negative deflection, and the duration of the wave (Mancuso et al., 2011; Gaja-Capdevila et al., 2021). Throughout the tests, pentobarbital (50 mg/kg i.p.) was used to anesthetize and the mice body temperature was maintained at a constant level using a thermostated heating pad. Electrophysiological tests were conducted at 8 (prior to the initiation of drug administration), 12, 14 and 16 weeks of age.,

### Histology

At the 16-week of age mice sacrificed with an overdose of pentobarbital and underwent transcardial perfusion with a 4% paraformaldehyde solution. Following perfusion, the lumbar segment of the spinal cord was carefully extracted, post-fixed for 2 hours, and subsequently cryopreserved in a solution of 30% sucrose in phosphate buffer (PB) at 4°C. Then, transverse 20-µm thick sections of the spinal cord were serially cut with a cryostat. For assessing the number of MNs, slices of L4-L5 spinal cord segments separated 100 μm were stained with a solution of 3.1mM cresyl violet for 3 h. MNs were identified by their anatomical localization in the ventral horn of the spinal cord and counted following strict criteria of their size (diameters lager than 20µm) and morphological characteristics (polygonal shape and prominent nucleoli).

### Neuromuscular junction assessment

After perfusion, the TA muscle was collected and underwent cryoprotection in PB 30% sucrose solution. To label neuromuscular junctions (NMJ), the muscle was sectioned in longitudinal slices of 50 µm thickness, collected in sequential series. Following a blocking step using 5% normal donkey serum, the sections were incubated for 48 hours at 4°C with primary antibodies, chicken anti-neurofilament 200 (NF200, 1:1000; AB5539, Millipore) and rabbit anti-synaptophysin (1:500; AB32127, Abcam). After washing steps, the sections were incubated overnight with secondary antibodies Alexa Fluor 594-conjugated secondary antibody (1:200; A11042-A21207, Invitrogen) and Alexa Fluor 488-conjugated α-bungarotoxin (1:500; B13422, Life Technologies). The prepared slides were then mounted using Fluoromount-G. Images were captured under confocal microscopy (LSM 700 Axio Observer, Carl Zeiss, 20x with a numerical aperture of 0.5), and maximum projection images were generated from z projections with a thickness of 1.3 µm. To evaluate the proportion of innervated NMJs, each individual endplate was classified as either occupied (when presynaptic terminals covered the endplate) or vacant (when there was no presynaptic label in contact with the endplate). A total of at least 60 endplates from four different fields were analyzed for each mouse.

### Statistical methods

Data are shown as mean ± SEM. Electrophysiological and functional measurements were analyzed with Two-way ANOVA, applying Bonferroni post-hoc test for multiple comparisons. Histological data were analyzed using One-way ANOVA with Bonferroni multiple comparisons test. The survival time of mice was evaluated using Kaplan-Meier analysis and Log-Rank test. Comparisons of means of gene and protein expression were analyzed by unrelated t-test. The computer-assisted analysis of the bands obtained in western blot was performed with AlphaEase FC software (Bonsai Technologies Group, S.A.). Statistical analysis was performed using SPSS (version 20, IBM, Armonk, NY, USA), and graphs were made using GraphPad Prism Software (version 5, La Jolla, CA, USA) and Microsoft Excel. Statistical significance was set at p<0.05.

## RESULTS

### Pharmacokinetics and target engagement of novel FKBP12 ligands

The newly designed 4-arylthiomethyl-1-carboxyalkyl-1,2,3-triazoles were synthesized with the aim of serving as potential ligands that can concurrently bind to both the ionic and hydrophobic pockets of the FKBP12/RyR1 complex (Aizpurua et al., 2021). These compounds demonstrated in vitro target engagement with FKBP12 through the rapamycin-inducible FKBP/FRP binding domain protein fragment complementation assay, thereby stabilizing cytosolic Ca^2+^ levels in human myotubes under nitro-oxidative stress (Aizpurua et al., 2021). Within this class of compounds, MP-001 (Fig. 1A) demonstrated superior binding affinity to FKBP12 and elicited co-localization of FKBP12/RyR1 in human myotubes under nitro-oxidative stress conditions (Aizpurua et al., 2021). MP-001 not only attenuated Ca^2+^ ER leakage through the RyR induced by nitro-oxidative stress but also demonstrated the capacity to rectify the heightened susceptibility to spontaneous Ca^2+^ release observed in conditions of ER Ca^2+^ overload resulting from the RyR2 R2474S mutation, which disrupts the interaction with the FKBP12.6 subunit (Lehnart et al., 2008). This manifestation is evidenced by the increased Ca^2+^ content released upon stimulation with caffeine (Suppl. Fig. 1).

**Figure 1.**
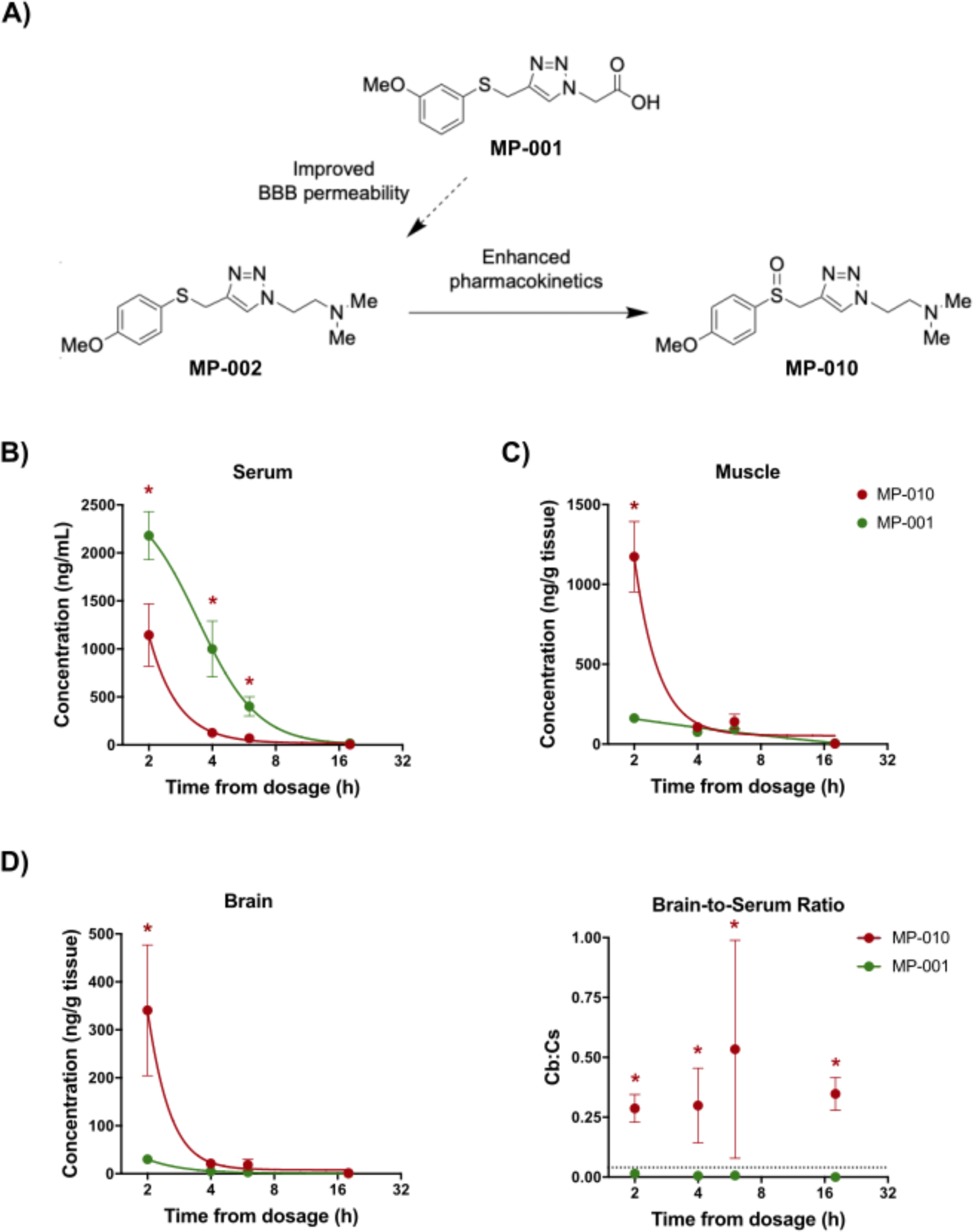
Chemical Structure of RyR Stabilizers and Concentration-Time Profiles in Mice Following a Single Oral Bolus of MP-010 or MP-001 at a Dose of 30 mg/kg. **A**) Chemical structures of FKBP12 ligands MP-001, MP-002 and MP-010. Compound MP-001 restored cytosolic calcium homeostasis in myotubes but show limited BBB permeability (See Fig. 1A-C). Modification to MP-002 improved BBB permeability, although the compound displayed too short serum half-time for systemic administration. Finally, the chemical incorporation of the sulfoxide function into MP-010 provided favorable pharmacokinetic parameters for development. **B,C,D**) Control mice were administered a single dose of MP-010 or MP-001 at 30 mg/kg by oral gavage, and their concentrations were assessed at 2, 4, 6, and 18 h post-dosing in the plasma (**B**), skeletal muscle (**C**), and brain (**D**). **D)** The calculated mean brain-to-serum (Cb:Cs) ratios were significantly higher for MP-010 than MP-001 and exhibited values above the threshold determining penetrability to the brain (Cb:Cs > 0.04, dashed line). The presented concentrations and values are the mean ± S.E.M. (n=3). *p< 0.05 vs. MP-001-treated mice.

Polar group exchange from carboxylic acid to N,N-dimethylamino moiety yielded compound MP-002 (Fig. 1A) with improved blood-brain-brain barrier (BBB) permeability. However, MP-002 exhibits very rapid metabolism, which results in low circulating levels, making it unsuitable for chronic treatment of the central nervous system (CNS). Concretely, MP-002 displayed a C_max_ of 5.1 ± 2.4 ng/mL with an inappreciable half-life in serum after an oral dosing of 30 mg/kg (data not shown).

Subsequently, for chronic treatment in the SOD1^G93A^ mouse model, we selected compound MP-010 (Fig. 1A) as an improved version of MP-002, incorporating a sulfoxide functional group to enhance its pharmacokinetic properties. MP-010 demonstrated favorable oral absorption, with a mean C_max_ of 1144.4 ± 397.2 ng/mL and a half-life of 3.0 h in serum (Fig. 1B), a C_max_ of 1173.1 ± 279.6 ng/g and a half-life of 2.5 h in muscle (Fig. 1C), and a C_max_ of 340.8 ± 166.9 ng/g and a half-life of 3.7 h in the brain after a dosing of 30 mg/kg (Fig. 1D). The brain-to-serum concentration ratio (Cb:Cs) of MP-010 was significantly greater than 0.04, even after 18 hours following dosing (Fig. 1D). This value is the threshold at which a molecule is commonly considered to be “brain penetrant,” as blood brain volume approximates 4% of total brain volume (Shaffer, 2010).

With respect to the stabilizing properties of RyR, both compounds MP-002 and MP-010 showed a complete rectification of the Ca^2+^ ER leak caused by the RyR2 R2474S mutation, as evidenced by the increased Ca^2+^ content released upon stimulation with caffeine (Suppl. Fig. 1).

Furthermore, we sought to evaluate another compound with in vivo stabilities comparable to those of MP-010 but with reduced or insignificant tissue distribution. Our aim was to determine that any positive effects of MP-010 are attributable to its mechanisms of action within the CNS. Based on this assumption, we chose MP-001 as an alternative compound, with a C_max_ and half-life of 2179.8 ± 305.9 ng/mL and 2.5 hours in serum (Fig. 1B), 163.0 ± 22.1 ng/g and 2.5 hours in muscle (Fig. 1C), and 30.3 ± 5.6 ng/g and 1.2 hours in the brain (Fig. 1D), respectively. Its Cb:Cs ratio was found to be less than 0.04 (Fig. 1D).

Therefore, we used MP-010 and MP-001 to investigate the hypothesis that modulating intracellular Ca^2+^ levels by facilitating the interaction between FKBP12 and RyR may confer neuroprotective effects in ALS.

### Evaluating the efficacy of compound MP-010 in alleviating neuromuscular deficits in SOD1^G93A^ mice

In the first phase of our study, we examined the potential of MP-010 to ameliorate motor impairments and neuromuscular dysfunction in SOD1^G93A^ transgenic mice. To this end, the animals were given two different doses of MP-010 (61 and 122 mg/kg) (Fig. 2A). Remarkably, the chronic administration of MP-010 at both doses examined did not result in any discernible differences in body weight between treated and vehicle groups of WT or SOD1^G93A^ mice (Fig. 2B). A notable increase in body weight was noted in the WT groups compared to the SOD1^G93A^ vehicle-treated group, particularly at the 12 to 16-week time points (Fig. 2B). This emphasizes the expected divergence in body weight trajectories between the WT and SOD1^G93A^ cohorts throughout the study.

**Figure 2.**
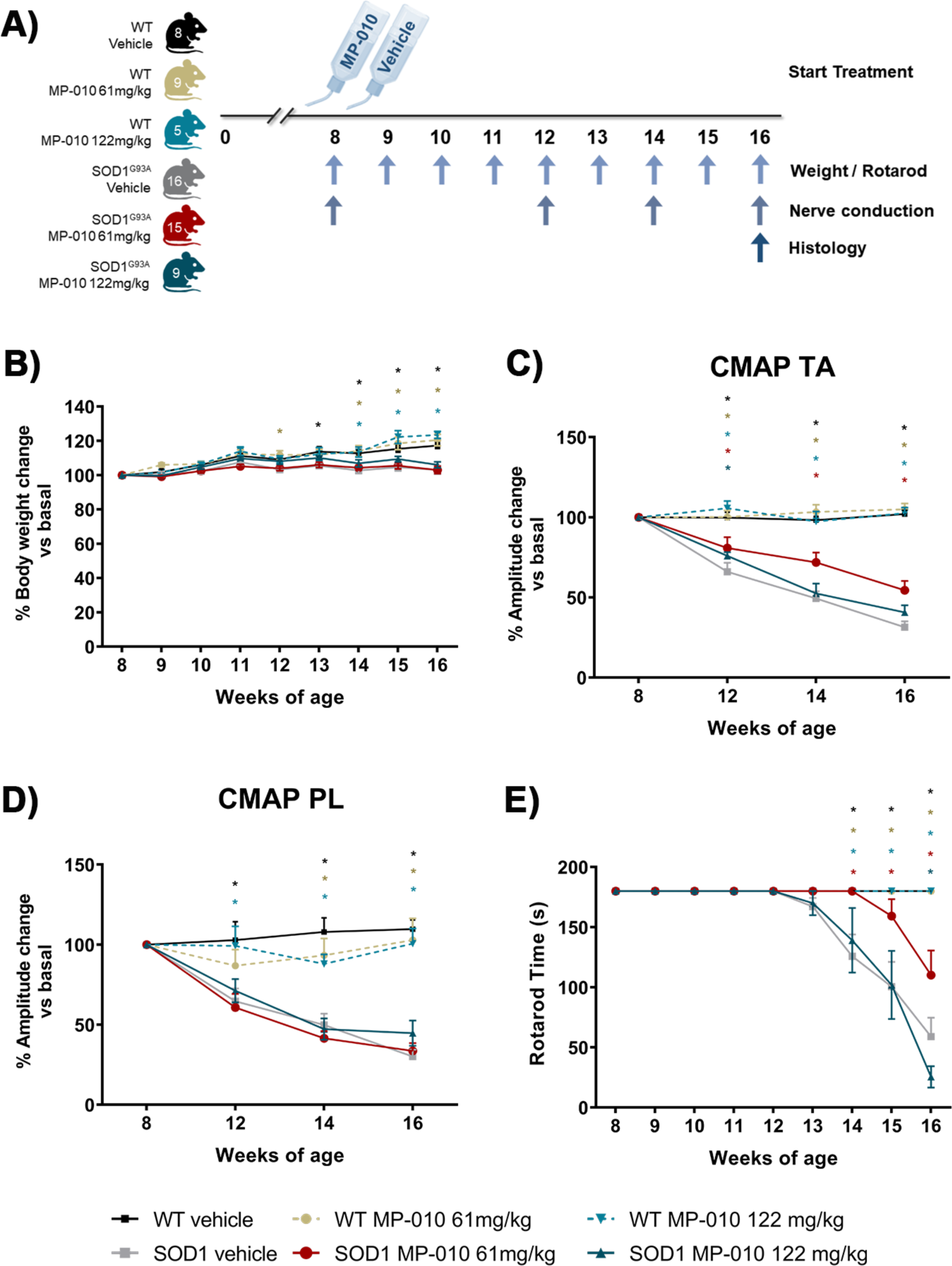
Improvement of Motor Function in SOD1^G93A^ Mice upon MP-010 Treatment. **A)** Schematic diagram of the experimental design for the trial whose data are shown in this figure and figure 3. **B)** Plot of the percentage of body weight for the different mice groups during the follow-up period. Body weight results are expressed as a percentage relative to baseline values for each mouse. **C-D)** Electrophysiological tests illustrating the preservation of compound muscle action potentials (CMAP) amplitude in the tibialis anterior (TA) muscle but not in the plantar interosseous (PL) muscle at 16 weeks of age. Amplitude values of CMAP are presented as a percentage relative to the baseline value for each mouse. **E)** Graph depicting the impact of different MP-010 treatments on functional outcome in the rotarod test. Mean ± SEM are represented in the plots. Two-way ANOVA with Bonferroni’s multiple comparisons test was employed. *p < 0.05 compared to the SOD1^G93A^ vehicle group. Each colored * represents the comparison between each group (defined by colors on panel A) and the SOD^G93A^ vehicle group (grey).

Electrophysiological assessments during the follow-up revealed that MP-010 treatment at a dose of 61 mg/kg prevented the decline of CMAP amplitude compared to untreated SOD1^G93A^ mice (Fig. 2C). Notably, at early (12 weeks), intermediate (14 weeks), and advanced (16 weeks) disease stages of the disease, MP-010 at 61 mg/kg exerted significant effects on CMAP amplitude preservation in the TA muscle of SOD1^G93A^ mice (Fig. 2C). In contrast, mice treated with the 122 mg/kg dose did not manifest a substantial improvement compared to the vehicle-treated group (Fig. 2C). In the PL muscle there were no significant improvements with any of the doses of MP-010 in the treated SOD1^G93A^ mice (Fig. 2D).

The rotarod test was used to assess locomotion and coordination, and to pinpoint the initiation of the symptomatic phase of the disease. The administration of MP-010 at the dose of 61 mg/kg yielded a significant improvement in rotarod performance at an advanced disease stage (14-16 weeks of age in SOD1^G93A^ mice) (Fig. 2E), delaying the onset of motor impairment by 2 weeks compared to untreated animals (13 weeks). In contrast, in accordance with the CMAP data, there was no change in rotarod performance between SOD1^G93A^ mice treated with 122 mg/kg and vehicle control. The beneficial effects of the 61 mg/kg regime were indicative of a deceleration in the progression of the disease, highlighting the potential therapeutic impact of MP-010 on mitigating motor deficits in SOD1^G93A^ mice.

At the end of the functional follow-up (16 weeks of age), a significant increase in the number of innervated NMJ within the TA muscle in SOD1^G93A^ mice receiving MP-010 treatment at both doses was noticed, compared to the vehicle-treated groups (Fig. 3A,B). This finding supports the observed preservation of CMAP amplitude in nerve conduction tests.

**Figure 3.**
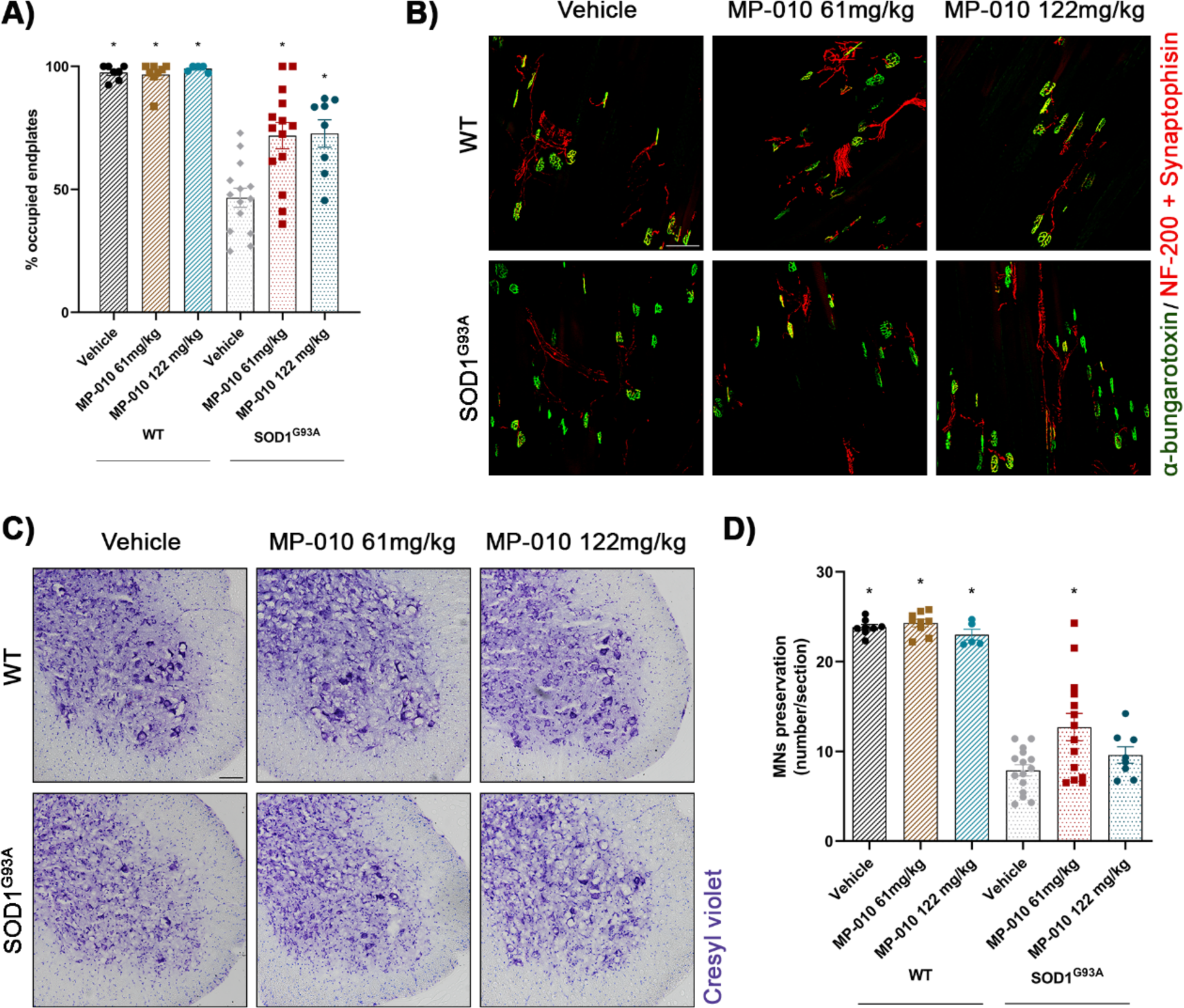
Preservation of NMJ Innervation and Spinal MNs in SOD1^G93A^ Mice upon MP-010 Treatment. **A)** Plot illustrating the percentage of neuromuscular junctions (NMJ) innervated (overlap of signals) in the different mouse groups. **B)** Representative confocal images of NMJ in tibialis anterior (TA) muscles of mice at 16 weeks of age, presented as maximum projections generated from 1.3 μm z projections. Immunolabeling against α-bungarotoxin (nicotine receptor, in green), neurofilament 200 + synaptophysin (synapse, in red) was performed to assess NMJ innervation. Scale bar: 100 µm. **C)** Representative images of lumbar spinal sections depicting MNs stained with cresyl violet from mice at 16 weeks of age. Scale bar: 100 μm. **D)** Plot showing the number of surviving spinal MNs (mean number of MNs per section ± SEM) in L4-L5 segments at 16 weeks of age in the different mouse groups. Mean ± SEM are represented in the plots. Statistical analysis employed one-way ANOVA with Bonferroni multiple comparisons test. *p < 0.05 compared to the SOD1^G93A^ vehicle group.

### Evaluating the efficacy of compound MP-010 in preserving MN loss in SOD1^G93A^ mice

At the end of functional follow-up, spinal cord was harvested for histological studies. The quantification of alpha MNs in the ventral horn of spinal cord sections stained with cresyl violet revealed that untreated control SOD1^G93A^ mice experienced a loss of over 60% of MNs compared to WT mice at 16 weeks of age (Fig. 3C,D). The administration of MP-010 at 61mg/kg significantly increased the number of surviving MNs in SOD1^G93A^ mice, whereas the dose of 122 mg/kg MP-010 did not mitigate MN loss (Fig. 3C,D).

### Evaluating the efficacy of compound MP-010 in lifespan in SOD1^G93A^ mice

After observing that MP-010 treatment confers motor and electrophysiological benefits, leading to reduced MN loss, a second trial in SOD1^G93A^ mice was conducted to investigate its potential to extend survival, particularly at the dosage of 61 mg/kg (Fig. 4A). Our observations revealed that MP-010-treated animals exhibited reduced weight loss compared to vehicle-treated controls, with significant differences becoming apparent at 18-19 weeks of age (Fig. 4B). The results obtained in the survival study indicated that MP-010 delayed mortality by approximately 7 days, extending the average lifespan from 132 ± 8.35 days in vehicle-treated SOD1^G93A^ mice to 139 ± 10.92 days in MP-010-treated mice (Fig. 4C).

**Figure 4.**
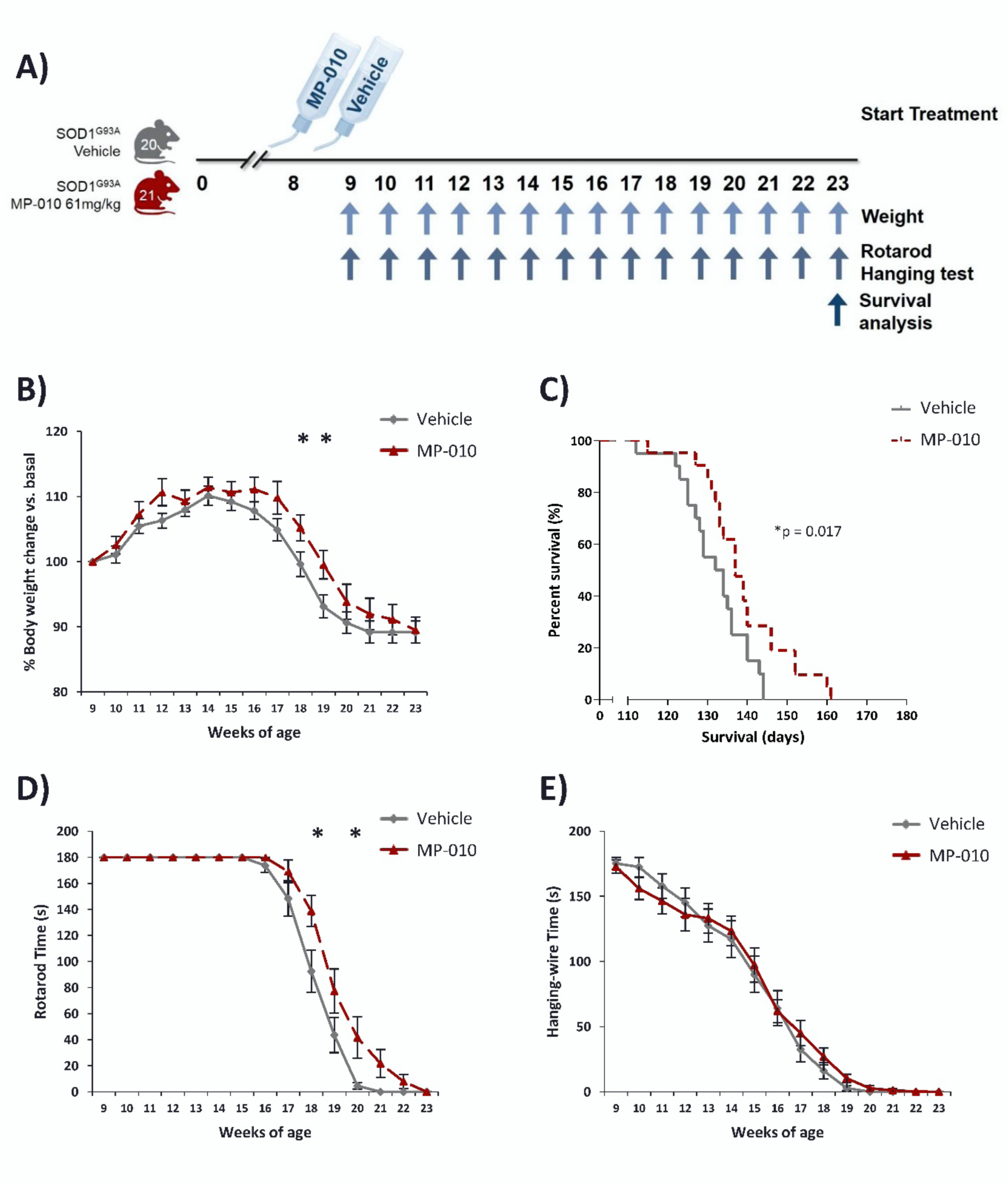
Enhanced Survival and Amelioration of Motor Symptoms in SOD1^G93A^ Mice Treated with MP-010. **A)** Schematic diagram of the experimental design of this trial. **B)** Plot illustrating the percentage of body weight for different mouse groups during the follow-up period. Body weight results are expressed as a percentage relative to baseline values for each mouse. **C)** Survival curves indicate that treated mice exhibited a longer lifespan compared to untreated counterparts, as determined by the Log-rank (Mantel-Cox) test. *p = 0.017 compared to the SOD1^G93A^ vehicle group. **D)** MP-010 treatment resulted in an improvement in motor coordination, as assessed by the rotarod test. **E)** MP-010 treatment did not lead to improved motor strength, as evaluated in the hanging-wire test. Mean ± SEM are represented in the plots. Two-way ANOVA with Bonferroni’s multiple comparisons test was employed. *p < 0.05 compared to the SOD1^G93A^ vehicle group.

Analyzing the outcomes of the behavioral tests, we again observed that MP-010-treated SOD1^G93A^ mice at the dose of 61 mg/kg demonstrated enhanced rotarod performance at advanced stages compared to vehicle-treated counterparts (Fig. 4D). Motor strength, assessed in the hanging-wire test, revealed no significant differences between the two groups of animals (Fig. 4E).

### Evaluating the efficacy of compound MP-001 in survival, motor function and muscular markers in SOD1^G93A^ mice

We also investigated on SOD1^G93A^ mice the therapeutic potential of the compound MP-001 (Fig. 5A), characterized by limited or negligible CNS availability (Fig. 1), thereby confining its mechanism of action to peripheral tissues. The body weight of all animals was monitored weekly from 8 weeks of age until 22 weeks. We observed reduced weight loss in the treated animals, with significant differences at the 13th and 15th weeks of age (Fig. 5B). However, as depicted in the survival analysis, MP-001 treatment did not significantly extend the survival of SOD1^G93A^ mice (Fig. 5C). Regarding the results obtained in the motor tests, no discernible improvement was observed in the SOD1^G93A^ mice treated with MP-001 (Figure 5D,E).

**Figure 5.**
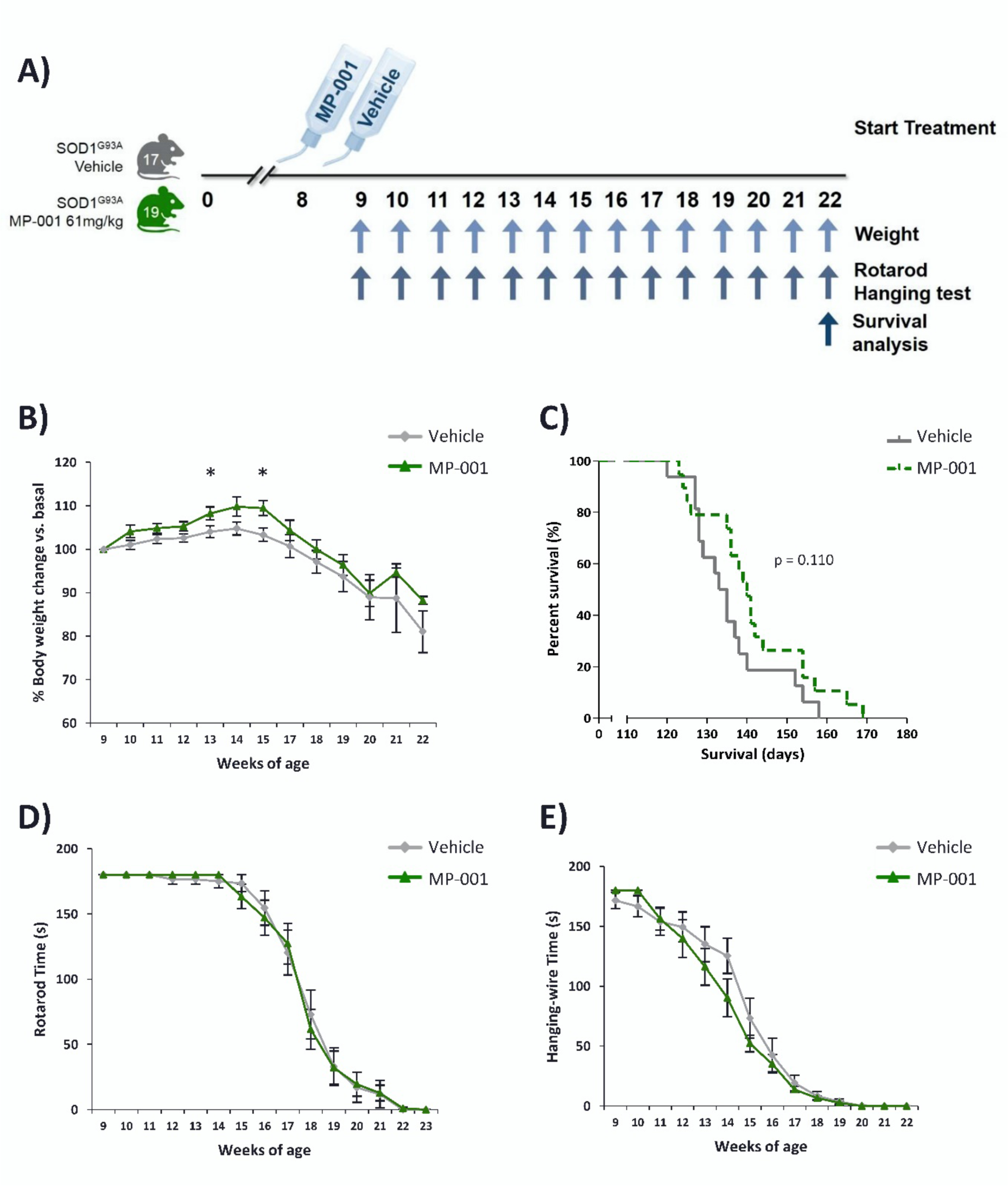
Marginal extension of survival and attenuation of weight loss in SOD1^G93A^ mice upon MP-001 administration. **A)** Schematic diagram of the experimental design for the trial to test MP-001. **B)** Mice treated with MP-001 exhibited reduced weight loss at 13-15 weeks of age. **C)** Treated mice displayed a slightly longer lifespan compared to untreated counterparts, but the differences were not statistically significant, as determined by the Log-rank (Mantel-Cox) Test. **D)** The treated and control animals demonstrated comparable results concerning motor coordination in the rotarod test. **E)** SOD1^G93A^ mice under treatment did not exhibit an improvement in motor strength, as assessed in the hanging-wire test. Mean ± SEM are represented in the plots. Two-way ANOVA with Bonferroni’s multiple comparisons test was employed. *p < 0.05 compared to the SOD1^G93A^ vehicle group.

## DISCUSSION

Our investigation centered on modulating RyR activity, specifically through a novel class of triazole molecules known as MP compounds that act as FKBP12 ligands (Aizpurua et al., 2021). The primary focus of our study was on MP-010, a RyR stabilizer with favorable pharmacokinetics and CNS permeability. Our results in SOD1^G93A^ mice, a widely accepted ALS model, demonstrated significant benefits following MP-010 administration. Notably, increased survival, protection against MN loss, muscle denervation, and enhanced neuromuscular function were observed, highlighting the potential of MP-010 as a therapeutic intervention for ALS. The robustness of our findings is emphasized by their replication in two distinct colonies of SOD1^G93A^ mice with a mixed hybrid B6SJL genetic background, across two independent laboratories. This replication is particularly noteworthy considering the potential genetic drift that hybrid ALS SOD1 colonies may undergo among different research labs, leading to variability in disease onset or symptom severity (Heiman-Patterson et al., 2005; Mancuso et al., 2012). This potential confounding factor makes the replication of our findings in two independent laboratories even more significant. Remarkably, despite differing ages of motor symptom onset between the two colonies (14 weeks in one laboratory and 17 weeks in the other), MP-010 demonstrated beneficial effects in both settings at the dose of 61 mg/kg. The compound exhibited remarkable tolerability and appeared devoid of apparent toxicity even at high doses. Doubling the dose did not induce a change in body weight, a recognized clinical symptom of pharmacological toxicity. Nevertheless, the compound’s therapeutic efficacy was compromised at this higher dose. This unexpected loss of efficacy could be attributed to off-target effects. These unintended interactions with non-target proteins could be interfering with the compound’s ability to reach its intended target and exert its therapeutic effects. Further investigations are warranted to identify and characterize specific off-target mechanisms, enabling the development of strategies to mitigate them and optimize the compound’s efficacy.

As previously introduced, cytosolic Ca^2+^ overload assumes particular importance in the vulnerability of spinal MNs subjected to ALS toxicity (Ho et al., 1996; Dekkers et al., 2004), and deficient Ca^2+^ handling from ER and mitochondrial stores has emerged as a key player in the pathogenesis of ALS. This sets the stage for a positive feedback loop that amplifies pathological Ca^2+^ overload, intensifying the neurodegenerative processes associated with the disease (Tadic et al., 2014; Leal and Gomes, 2015). Mutations in genes such as *SIGMA1R* and *VAPB*, identified in ALS patients, underscore the significance of ER and mitochondrial proteins in Ca^2+^ homeostasis (De vos et al., 2012; Bernard-Marissal et al., 2015). In the context of the present study, the rationale for targeting RyR stemmed from its central role in Ca^2+^-induced Ca^2+^ release from the ER, aiming to prevent excessive cytosolic Ca^2+^ accumulation, especially in the most vulnerable MNs associated with ALS. The outcomes of the current study, demonstrating the therapeutic benefits of a drug promoting the interaction between RyR and its endogenous ligand FKBP12 in the SOD1^G93A^ mouse model of ALS, are in accordance with earlier research that has emphasized reductions in FKBP12 in the context of ALS (Kihira et al., 2005). Also in the SOD1^G93A^ mouse, various alternative strategies to reduce the Ca^2+^ burden on MNs have been successfully implemented. These strategies include the application of the AMPA receptor antagonist talampanel (Paizs et al., 2011) and SIGMA1R agonists (Mancuso et al., 2011; Gaja-Capdevila et al., 2021), both exerting their effects through the modulation of extracellular Ca^2+^ influx to cytosol.

The dysregulation of RyR signaling emerges as a fundamental contributor to the pathogenesis of various neurodegenerative disorders, notably Alzheimer’s disease (AD). Specifically, RyRs have been demonstrated to interact with amyloid-beta (Aβ) peptide, a protein known to play a pivotal role in AD development. The binding of Aβ to RyRs can modulate their function, resulting in heightened Ca^2+^ release from the ER, subsequently leading to excitotoxicity, neuronal death, and cognitive deficits (Liu et al., 2012; Lacampagne et al., 2017; Yao et al., 2020, 2022). Although our investigation cannot definitively ascertain the existence of Ca^2+^ leakage through RyR in ALS MNs or whether this pathological phenomenon is replicated in the SOD1^G93A^ mouse model, the administration of the compound MP-010 emphasizes the potential therapeutic significance of targeting FKBP12 to safeguard the RyR-dependent proper release of Ca^2+^ and alleviate the downstream effects of disrupted Ca2+ homeostasis. This approach offers a targeted intervention for ALS. However, while it is important to note that an aberrant increase in RyR-mediated Ca^2+^ release can contribute to disrupting neuronal homeostasis, triggering excitotoxicity, conversely, diminished RyR function may compromise the crucial Ca^2+^ signaling required for proper neuronal operation. This impairment might lead to a breakdown in synaptic transmission and plasticity, potentially contributing to the manifestation of motor deficits in individuals with ALS. Consequently, inhibiting RyR under this premise may not be a plausible strategy for treating neurons susceptible to ALS. Indeed, prior research utilizing dantrolene, an RyR1 inhibitor clinically used to treat malignant hyperthermia, failed to demonstrate significant improvements when administered to the SOD1^G93A^ mouse model (Staats et al., 2012), confirming the inadequacy of RyR inhibition as a therapeutic avenue. In contrast, compounds exhibiting RyR-stabilizing activity, previously recognized under the nomenclature “rycals” (Fauconnier et al., 2010), adopt a distinct approach by modulating RyR channel opening. These compounds achieve this modulation by facilitating the interaction between the topoisomerase FKBP12 and RyR, particularly in pathological scenarios characterized by persistent Ca^2+^ leakage (Fauconnier et al., 2010). Thus, the strategic targeting of RyR modulation through rycal compounds offers a more nuanced and potentially effective therapeutic strategy compared to direct inhibition.

Comparative analysis with MP-001, another compound characterized by limited CNS availability, provides additional insights. The lack of MP-001 effect underscore the significance of targeting Ca^2+^ homeostasis at the CNS level to achieve therapeutic efficacy. Thus, it dismisses the possibility that the therapeutic effects could be mediated by the modulation of Ca^2+^ dynamics in cells of the innate immune system. It is crucial to consider this aspect, especially given that immune cells heavily rely on Ca^2+^-dependent processes and tightly regulated Ca^2+^ homeostasis (Peters and Raghavan, 2011), and that the innate immune response appears to play an active role in the pathophysiology of ALS and the SOD1G93A mouse model (Nguyen et al., 2004; Komine et al., 2018).

While our study presents promising results, several avenues for future research emerge. Elucidating the molecular mechanisms underlying the observed effects of MP-010, particularly its impact on Ca^2+^ dynamics and neuronal survival, is critical for a comprehensive understanding of ALS pathology. Additionally, considerations of toxicity, dosage optimization, and potential side effects are paramount before translating these findings into clinical trials. In conclusion, our study contributes to the evolving landscape of ALS research by highlighting the therapeutic potential of targeting intracellular Ca^2+^ dynamics through the modulation of RyR activity. MP-010, with its favorable preclinical outcomes, offers a promising avenue for further exploration and potential clinical development.

## AUTHOR CONTRIBUTION

X.N, R.O., A.L.M., A.V-I., F.J.G-B., J.F. and A.A-A. contributed to design the study. J.I.M., J.M.A., and M.S-A. designed and synthesized the compounds. L.M-M., N.G-C., L.M. and M.H-G. performed experiments and analyzed data. X.N, R.O., A.L.M., A.V-I., F.J.G-B, L.M-M., N.G-C., L.M. contributed to the preparation of the manuscript. X.N, R.O., A.L.M., and F.J.G-B. supervised and coordinated all the work.

## ACKNOWLEDGEMENTS

Dr. Takashi Sakurai is gratefully acknowledged for his generous gift of HEK293 cells stably and inducibly expressing mutant and WT forms of RyR2 and co-expressing R-CEPIA1er, which were instrumental in the successful completion of this work. The authors thank for technical and human support provided by Mª Carmen Sampedro from SGIker of UPV/EHU and European funding (ERDF and ESF), and by Neus Hernández, Mónica Espejo and Jessica Jaramillo from the UAB for their excellent technical support.

