## Supplementary Figure 1 for "Novel FKBP12 ligand promotes functional improvement in SOD1^G93A^ ALS mice"

### SUPPLEMENTARY MATERIAL

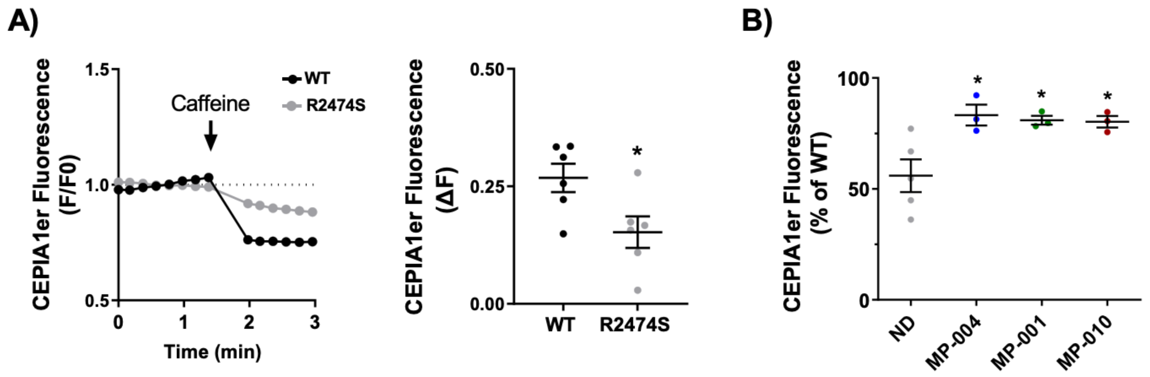


**Supplementary Figure 1.**

**A)** Caffeine-induced ER $Ca^{2+}$ release is significantly reduced in HEK293 cells expressing the mutant R2474S-RyR2 compared to WT cells. This suggests that $Ca^{2+}$ stores in the ER are depleted due to the continuous leakage of mutant RyR2 channels. *p<0.05 (unpaired t-test).

**B**) Pretreatment with MP compounds increase caffeine-induced ER $Ca^{2+}$ release in RyR2 HEK mutants, suggesting that RyR2 leakage is mitigated. Cells were pretreated or not for 1 h with MP compounds. 10 mM caffeine was used to empty ER stores. ND (non-drug) refers to the effect of caffeine in non-treated HEK293 expressing RyR2 R2474S. Assays were performed in triplicates. *p<0.05, One-Way ANOVA post-hoc Dunnet's multiple comparisons test vs ND.
